# Endurance Exercise Elevates Plasma Mature Brain-Derived Neurotrophic Factor through Child- and Adulthood, with Amplified Effects in Adults

**DOI:** 10.1101/2024.11.06.622216

**Authors:** Sebastian Edman, Julia Starck, Linnéa Corell, William Hangasjärvi, Amelie von Finckenstein, Mikael Reimeringer, Stefan Reitzner, Jessica Norrbom, Marcus Moberg, Ferdinand von Walden

## Abstract

**Background:** Brain-derived neurotrophic factor (BDNF) is a neurotrophin that plays a central role in neuronal health. BDNF exists in two primary isoforms, the mature form (mBDNF) and its precursor (proBDNF), with opposing downstream effects on neuronal function. The positive effect of exercise on plasma levels of the BDNF-isoforms has been extensively studied in adults. However, equivalent investigations are lacking in children and youth.

**Methods:** Twenty healthy children (9-12 years old), 19 adolescents (13-17 years old), and 39 adults (23-49 years old) donated venous blood before and after a 45-minute run. Platelet-poor plasma was analyzed for pro– and mBDNF using enzyme-linked immunosorbent assay. Maximal oxygen uptake and anthropometric data were assessed in all participants, while Tanner stage, circulating sex hormones, and accelerometry-based activity level were assessed in children and adolescents.

**Results:** We found that children, adolescents, and adults have similar basal levels of pro-(337-543 pg × ml^−1^) and mBDNF (650-1111 pg × ml^−1^). For children and adolescents, basal levels of mBDNF correlated with average time spent in vigorous activity (r=0.5794, p=0.0015). In response to acute endurance exercise, mBDNF increased by 701 ± 946, 1232 ± 1105, and 1557 ± 1168 pg × ml^−1^ in children, adolescents, and adults, respectively. The acute endurance exercise did not affect proBDNF levels (p>0.05).

**Conclusion:** Our results demonstrate that basal plasma pro– and mBDNF levels do not depend on age and maturity. Plasma mBDNF levels increase following endurance exercise in all age groups, but to a greater extent in adults.

**Highlights/Impact:**

- We show that in children and adolescents, regular vigorous physical activity is key to increased basal levels of plasma mBDNF, a factor linked to neuroplasticity and brain health.
- The ability to elevate mBDNF through exercise is present across all age groups, with the greatest increase in adults.
- The mBDNF response to physical exercise seems to be independent of underlying physical fitness.
- Our findings suggest that basal plasma mBDNF levels may reflect the cumulative effects of repeated exercise rather than an individual’s overall physical fitness.

## Introduction

Brain-derived neurotrophic factor (BDNF) is a neurotrophin that plays a central role in brain health throughout life. It influences key processes such as neuronal differentiation, synapse function, and cognitive development^1^. BDNF exists in two primary forms: the mature BDNF (mBDNF), which promotes neuronal survival and plasticity through its interaction with the TrkB receptor^2^, and its precursor, proBDNF, which binds to the p75NTR receptor, inducing apoptosis^3,4^. The balance between these isoforms is crucial for maintaining proper neural function, with their opposing actions often described as a “yin and yang” dynamic that regulates neuronal survival and function^2^. However, due to the cleavage capacity of proBDNF to mBDNF^4^, the high levels of proBDNF observed within skeletal muscle could also be regarded as a precursor pool for circulating mBDNF^5^. In the circulation, the BDNF isoforms can be found either bound to platelets or freely circulating in plasma^6^. While serum levels capture the total BDNF (both bound and unbound), plasma measures specifically reflect the free, unbound fraction. The free BDNF is thought to cross the blood-brain barrier bidirectionally^7^, providing a link between peripheral and central BDNF levels and contributing to both central and peripheral neuroprotection.

Physical exercise has repeatedly been reported to have neuroprotective effects, lowering the rate of cognitive decline^8^, and increasing mental health status^9^, as well as memory, attention, and learning ability^10^. The cognitive and neuroprotective benefits associated with exercise are oftentimes linked to the exercise-mediated increases in cerebral BDNF levels of mice^11–13^ and the subsequent release of BDNF into the circulation^14,15^. Acute exercise is a potent stimulator of both plasma and serum BDNF^16^, however, the response can vary based on several factors, such as exercise duration^17^ and intensity^18^. It has further been suggested that the plasma BDNF release in response to acute exercise is ameliorated by prior exercise training^19^ or physical fitness^20^. Long-term training interventions have also been associated with higher basal levels of plasma BDNF^19^, however, these long-term training effects are less consistent across the literature than that of the transient BDNF release into circulation with acute exercise^21,22^.

The effects of exercise on BDNF levels in children have, to date, been investigated only in a small number of publications, unlike the effect in adults, leading to some uncertainty in the literature. While a few studies have reported a positive association between physical activity levels and basal plasma BDNF in children^23^, others have found no such correlation^24^. Additionally, there is evidence suggesting a relationship between physical fitness levels and higher plasma BDNF in children^25^. During postnatal development, BDNF levels in the central nervous system (CNS) are thought to increase significantly^26^, with BDNF mRNA levels rising from infancy and plateauing in early adulthood, as observed in human postmortem samples^27^. However, studies in other species, such as monkeys^28^ and rats^29^, have shown opposing trends in BDNF mRNA expression during development and aging. These discrepancies make it challenging to draw definitive conclusions about BDNF regulation across childhood and adolescence. Mechanistically, sex hormones, particularly estrogen, are believed to significantly influence BDNF levels in the CNS and circulation^30,31^. Estrogen has been shown to regulate BDNF gene and protein expression^32,33^, which may explain potential maturity-related differences in BDNF.

Therefore, the aim of the present study is to determine the basal concentrations of proBDNF and mBDNF in platelet-poor plasma among healthy children, adolescents, and adults, as well as to investigate the acute effects of intense endurance exercise on these levels. Additionally, we aim to explore whether baseline levels or exercise-induced changes in BDNF are influenced by factors such as physical fitness, weekly physical activity, developmental stage, and circulating sex hormones.

## Methods

### Ethical approval

The current project received ethical approval from the Swedish Ethical Review Authority 2023-06-22 (Dnr 2023-02343-01) and 2023-12-19 (Dnr 2023-07692-02). The study was conducted in line with the principles of the Declaration of Helsinki.

### Participants

Seventy-eight individuals aged 9 to 49 (40 women, 38 men) were recruited to the study (Table 1), of which 20 were children (9-12 yo), 19 were adolescents (13-17 yo) and 39 were adults (18-49 yo). Before inclusion, each participant filled out a health questionnaire to ensure all participants could perform the tests without risk of complications. Exclusion criteria included inability to perform the tests correctly, neuromuscular or known blood-borne diseases, injuries or other diseases that make endurance– and strength training inappropriate, or conditions or current treatments that can impact the training effect.

**Table 1.** Participant characteristics.

|  | Women | Men | Total |
| --- | --- | --- | --- |
| **Children aged 9-12yrs** | **N=10** | **N=10** | **N=20** |
| Age (years) | 10.4 ± 1.0 | 10.9 ± 0.9 | 10.6 ± 0.9 |
| Weight (kg) | 40.5 ± 8.5 | 40.7 ± 6.8 | 40.6 ± 7.5 |
| Height (cm) | 150.1 ± 15.3 | 148.9 ± 6.7 | 149.5 ± 11.5 |
| Body fat (%) | 20.2 ± 7.0 | 17.6 ± 5.2 | 18.9 ± 6.1 |
| VO<sub>2</sub>max (ml/kg/min) | 45.2 ± 4.2 | 51.6 ± 5.5 | 48.6 ± 5.8 |
| Tanner stage - breast/genitalia | 2.1 ± 0.9 | 2.3 ± 0.9 | 2.2 ± 0.9 |
| Tanner stage – body hair | 1.9 ± 0.9 | 1.8 ± 0.6 | 1.9 ± 0.7 |
| **Adolescents aged 13-17yrs** | **N=8** | **N=11** | **N=19** |
| Age (years) | 14.5 ± 1.5 | 13.9 ± 1.0 | 14.2 ± 1.2 |
| Weight (kg) | 58.1 ± 6.3 | 56.6 ± 8.5 | 57.2 ± 7.5 |
| Height (cm) | 169.0 ± 7.2 | 169.2 ± 9.5 | 169.1 ± 8.4 |
| Body fat (%) | 22.5 ± 3.4 | 14.8 ± 7.0 | 18.1 ± 6.8 |
| VO<sub>2</sub>max (ml/kg/min) | 43.6 ± 3.8 | 52.3 ± 6.0 | 49.2 ± 6.7 |
| Tanner stage - breast/genitalia | 4 ± 0.8 | 3.3 ± 0.7 | 3.6 ± 0.8 |
| Tanner stage – body hair | 3.9 ± 0.7 | 3.6 ± 0.9 | 3.6 ± 0.9 |
| **Adults** | **N=22** | **N=17** | **N=39** |
| Age (years) | 40.0 ± 8.4 | 41.3 ± 6.4 | 40.5 ± 7.5 |
| Weight (kg) | 63.8 ± 6.7 | 85.0 ± 14.8 | 73.0 ± 15.2 |
| Height (cm) | 166.9 ± 6.5 | 183.3 ± 5.7 | 174.1 ± 10.2 |
| Body fat (%) | 23.8 ± 6.2 | 19.8 ± 5.7 | 22.0 ± 6.2 |
| VO<sub>2</sub>max (ml/kg/min) | 42.8 ± 6.2 | 44.6 ± 6.9 | 43.5 ± 6.5 |

VO<sub>2</sub>max = Maximal oxygen uptake. Values are mean ± SD.

### Overall study design

Following enrollment in the study, children, and adolescents performed a cycle ergometer test (Ekblom-Bak), and all participants performed a maximal oxygen uptake test. Next, participants were scheduled for the acute exercise intervention on a separate occasion.

Lactate was measured during the run, and pro– and mBDNF before and after. Resting samples from all children and adolescents were evaluated for blood testosterone, sex hormone-binding globulin, follicle-stimulating hormone, luteal hormone, and estradiol. Sex hormones were analyzed by Karolinska University Laboratory. Children and adolescents also conducted a Tanner stage self-assessment during this session. Following the acute exercise intervention, children and adolescents were also instructed to start wearing physical activity monitors after a few days of rest.

### Cycle ergometer test

A submaximal cycle ergometer test, the Ekblom-Bak test, was performed on a Monark 828E ergometer (Monark, Vansbro, Sweden) with seat and handle height adjusted individually. The participants were equipped with a chest strap heart rate monitor (smartLAB, Heidelberg, Germany) and introduced to the Borg Rating of Perceived Exertion (RPE) scale. The test followed a standard protocol, and maximal oxygen uptake (VO_2_max) was estimated using the updated Ekblom-Bak equation^34^.

### Maximal oxygen uptake test

Maximal oxygen uptake (VO_2_max) was measured during a maximal treadmill running test, using a Rodby RL2500E treadmill (Vänge, Sweden). Continuous breath-by-breath gas exchange assessment was measured throughout the test, using a COSMED Quark CPET with Omnia software (Rome, Italy). Participants wore a chest strap heart rate monitor (smartLAB) for continuous heart rate monitoring. After a few minutes of warm-up, the test was started and the treadmill was set to 8-10 km/h, depending on the capacity of the individual. Each minute the treadmill velocity was increased by 0.5-1 km/h until the participant reached their highest tolerable running speed without significant form breakdown, after which the test continued with increasing the inclination by 2% for one minute, followed by 1% every following minute. The test proceeded until exhaustion and the participant was asked to rate exertion on the Borg RPE scale at the end of the test. The maximal oxygen uptake test lasted on average 09:06 ± 01:50 (Min:Sec), with the test length not significantly differing between the groups (08:58 ± 02:12, 08:50 ± 02:01, and 09:17 ± 01:32 for children, adolescents, and adults, respectively). The highest VO_2_ average for 45 seconds was considered as the participant’s VO_2_max. The VO_2_max value was finally divided by body weight and presented as ml O_2_/kg/min.

### Exercise intervention – 45 min running

On the morning of the exercise intervention, all participants received a standardized breakfast, consisting of 5 ml x kg bodyweight of the nutrition drink Resource Komplett (21 / 5 / 5.6 g of carbohydrates / fat / protein per 100 g drink). The drink was consumed one hour before starting the running test. Upon arrival, each participant received a peripheral venous catheter (PVC) in the antecubital vein on the forearm. Children and adolescents were instructed to apply local anesthetic cream (EMLA; Aspen Pharma TL, Dublin, Ireland) 1-3 hours in advance to reduce discomfort and pain. During the 45-minute run, the participants were equipped with a chest strap heart rate monitor (Garmin, Olathe, KS, USA) and were instructed to run at a steady pace while trying to run as far as possible on a 200-meter indoor running track during the 45-minute period. Heart rate (HR) data was collected as average HR for one whole minute every 10 min and immediately after the run. A weighted mean value of these HR-data points was divided by max HR (obtained either during the VO2-max test or the 45-minute run). Borg RPE scale was also assessed at 10-, 20-, 30-, 40-, and 45 min of running. Blood samples were collected before starting the exercise, 45 min after the standardized breakfast and immediately after the running, in both heparinized tubes and 1 ml syringes. Blood samples from the syringe were immediately analyzed for lactate levels using a blood gas analyzer (ABL800 Flex®, Radiometer Copenhagen, Denmark).

### Enzyme-linked immunosorbent assay

Immediately following collection in a heparinized tube, blood was centrifuged at 3000 g in 4°C for 10 min. The upper 75 % of the plasma was collected from the tube and subsequently centrifuged again. The upper 75 % was collected once more and immediately frozen on dry ice. The platelet-poor plasma obtained before and immediately after running were subsequently analyzed for pro– and mBDNF levels using Enzyme-linked immunosorbent assay (ELISA) according to the manufacturer’s instructions (Human BDNF Quantikine Immunoassay 8DBD00, R&D Systems and Human Pro-BDNF ELISA, #BEK-2237, Biosensis). In total, platelet-poor plasma from 19 children, 17 adolescents, and 34 adults was used for proBDNF measurements, and from 19 children, 19 adolescents, and 34 adults for mBDNF measurements.

### Physical activity assessment

Physical activity was measured in all children and adolescents for one week. The children were asked to wear accelerometers (ActiGraph wGt3x-BT, Actigraph LLC; Pensacola FL) on the waist and reported wear time and daily activities. The data was analyzed with R package GGIR version 3.1-4^35^. A day was considered valid if there was data collected for at least 10 hours. Participants with less than two valid days were excluded from analysis of the accelerometry data, resulting in data from participants 17 children and 10 adolescents being analyzed. They wore the accelerometer for an average of 5 ± 2-6 valid days (median ± range). Physical activity was measured as time spent in light, moderate, and vigorous activity. Cut-off for light activity was set to 63.3 for ages 9-12 and 26.9 for ages 13-17; for moderate activity to 142.6 for ages 9-12 and 332.0 for ages 13-17; and for vigorous activity to 464.6 for age 9-12 and 558.3 for age 13-17, all expressed in gravitational units^36–38^.

### Statistics

Comparisons of basal and delta mBDNF and proBDNF levels in children, adolescents, and adults were performed using a one-way ANOVA with Tukey post hoc testing to correct for multiple comparisons. A two-way repeated measures ANOVA was also conducted on the pre-and post-values of mBDNF and proBDNF of all groups to assess the effect of exercise in each group individually. Prior to analysis using ANOVA, mBDNF, and proBDNF were log-transformed to better meet the normality assumption assessed by the Shapiro-Wilks test.

Correlations were calculated using Pearson’s R, except for non-continuous data (Borg RPE scale, and Relative HR max), in which case Spearman’s rank coefficient was used. Data is presented as mean ± SD unless otherwise specified. The level of significance was set at *p* < 0.05. Statistical analysis was conducted using GraphPad Prism 10 for Mac.

## Results

In the rested state, 45 minutes after the standardized breakfast, plasma levels of proBDNF (337 ± 445 – 543 ± 675 pg × ml^−1^) and mBDNF (650 ± 605 – 1111 ± 1022 pg × ml^−1^) were not different between children, adolescents, and adults (Figures 1A & B). To further assess whether maturity, rather than age, affected basal pro– and mBDNF levels, we stratified the data from the children and adolescents based on puberty stage and sex hormone levels. We found no effect of maturity using Tanner stage (Genitalia/Breast & Hair) assessment of either BDNF isoforms at rest (Supplemental Figures 1A-B & 1D-E). Likewise, sex hormone concentrations for children and adolescents were not associated with basal levels of pro-or mBDNF (Supplemental Figures 1C-F), except for follicle-stimulating hormone levels (S-FSH) showing a moderate correlation with plasma mBDNF levels in girls (Pearsons r = 0.471, p<0.05; Supplemental Figures 1F). Furthermore, irrespective of age or maturity, resting levels of mBDNF were not associated with fitness levels assessed by either incremental treadmill running (Figure 1C) or estimated using the Ekblom-Bak ergometer test (Children and adolescents only; Figure 1D). By contrast, basal levels of circulating mBDNF were instead associated with physical activity levels, assessed with accelerometers in children and adolescents only (Figures 1E-H). Specifically, vigorous activity showed a moderate (r=0.5794, p=0.0015) correlation to basal levels of mBDNF (Figure 1H). Lower intensity activity levels, i.e. moderate-, and light activity as well as inactivity, showed no association with basal mBDNF (Figure 1E-G). Furthermore, for children and adolescents, time spent in vigorous physical activity (Figure 1L), but not moderate-to-light physical activity or inactivity (Figure 1I-K), was associated with VO_2_max levels.

**Figure 1.**
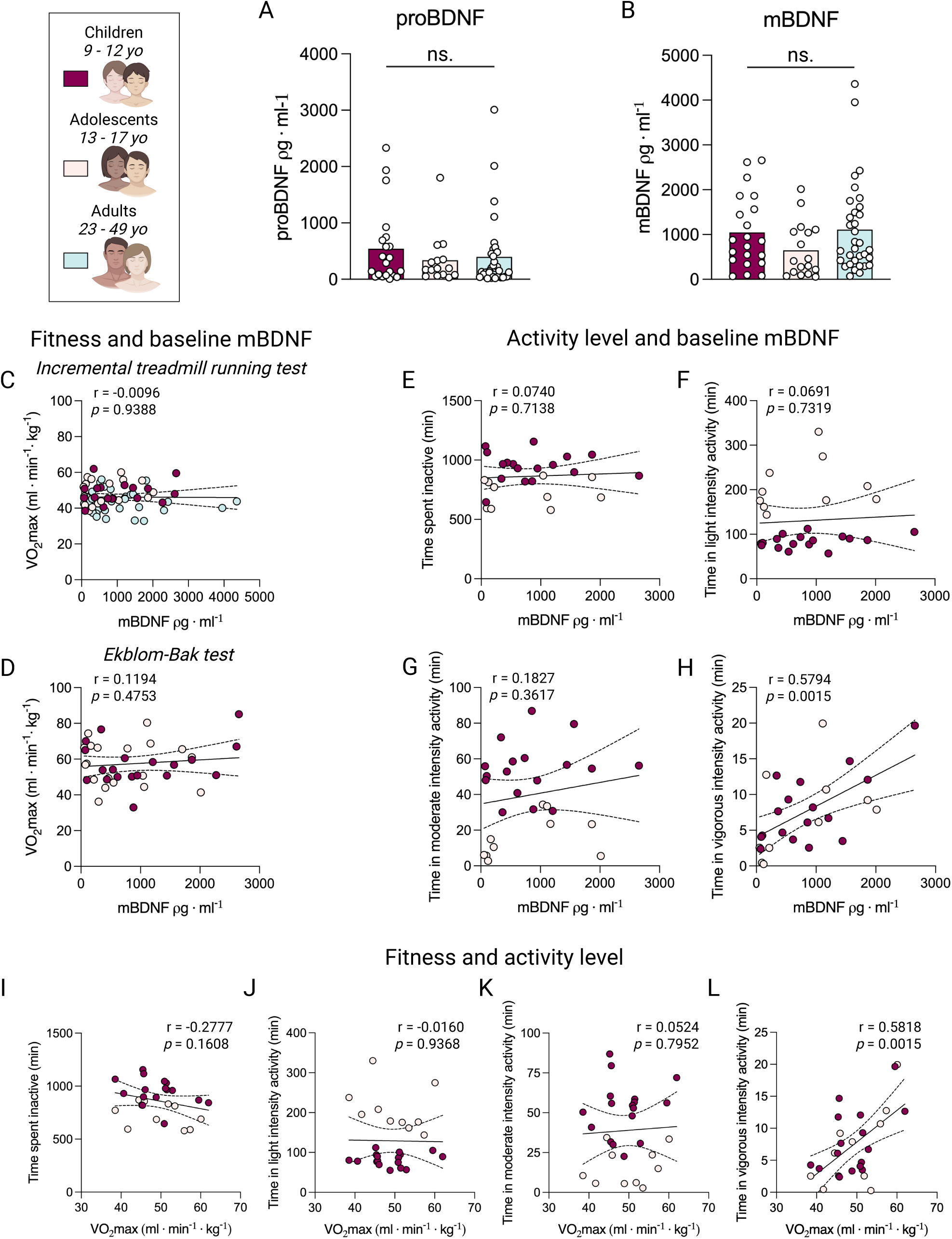
Plasma pro– and mBDNF at rest in children, adolescents, and adults. **A-B**: proBDNF levels (A) and, mBDNF levels (B) in plasma at rest. ns. = not significant. Group differences were analyzed by One-way ANOVA. **C-D**: Correlation of plasma mBDNF levels at rest with VO_2_max measured during incremental treadmill running (C) and estimated VO_2_max using the Ekblom-Bak test in children and adolescents (D). **E-H**: Correlation of plasma mBDNF levels at rest with time spent inactive (E), in light activity (F), in moderate-intensity activity (G), and in vigorous-intensity activity (H) during the day in children and adolescents. **I-L**: Correlations of VO_2_max measured during incremental treadmill running with time spent inactive (I), in light activity (J), in moderate-intensity activity (K), and in vigorous-intensity activity (L) during the day in children and adolescents. C-L was analyzed using Pearson correlation. VO_2_max = Maximal oxygen uptake

Following the 45-minute run, we observed no change in plasma proBDNF levels (Figure 2A), with the delta values of proBDNF showing no association with maturity (Supplemental Figures 2A-C). However, mBDNF concentrations increased in all age groups following exercise (701 ± 946 – 1557 ± 1168 pg × ml^−1^, p<0.05; Figure 2B). The exercise effect was greater in adults (140%) as compared to children (67%, p<0.05 for Children vs Adults; Figure 2B), suggesting an effect of age. The participant’s body weight showed a moderate correlation with the exercise-mediated mBDNF release (r=0.3827, p=0.001). We further observed that children and adolescents in the more advanced Tanner stages (4 & 5) had a significantly larger effect of exercise than children in Tanner stages 2 & 3 (p<0.05 for Tanner assessment using Hair; Supplemental Figures 2D-E). Moreover, circulating testosterone levels tended to correlate moderately with delta mBDNF levels in boys primarily (r=0.397, p=0.083, Supplemental Figure 2F). Neither underlying fitness (Figures 2C-D) nor activity level (Figures 2E-H) were associated with the plasma mBDNF change elicited by the 45-minute run.

**Figure 2.**
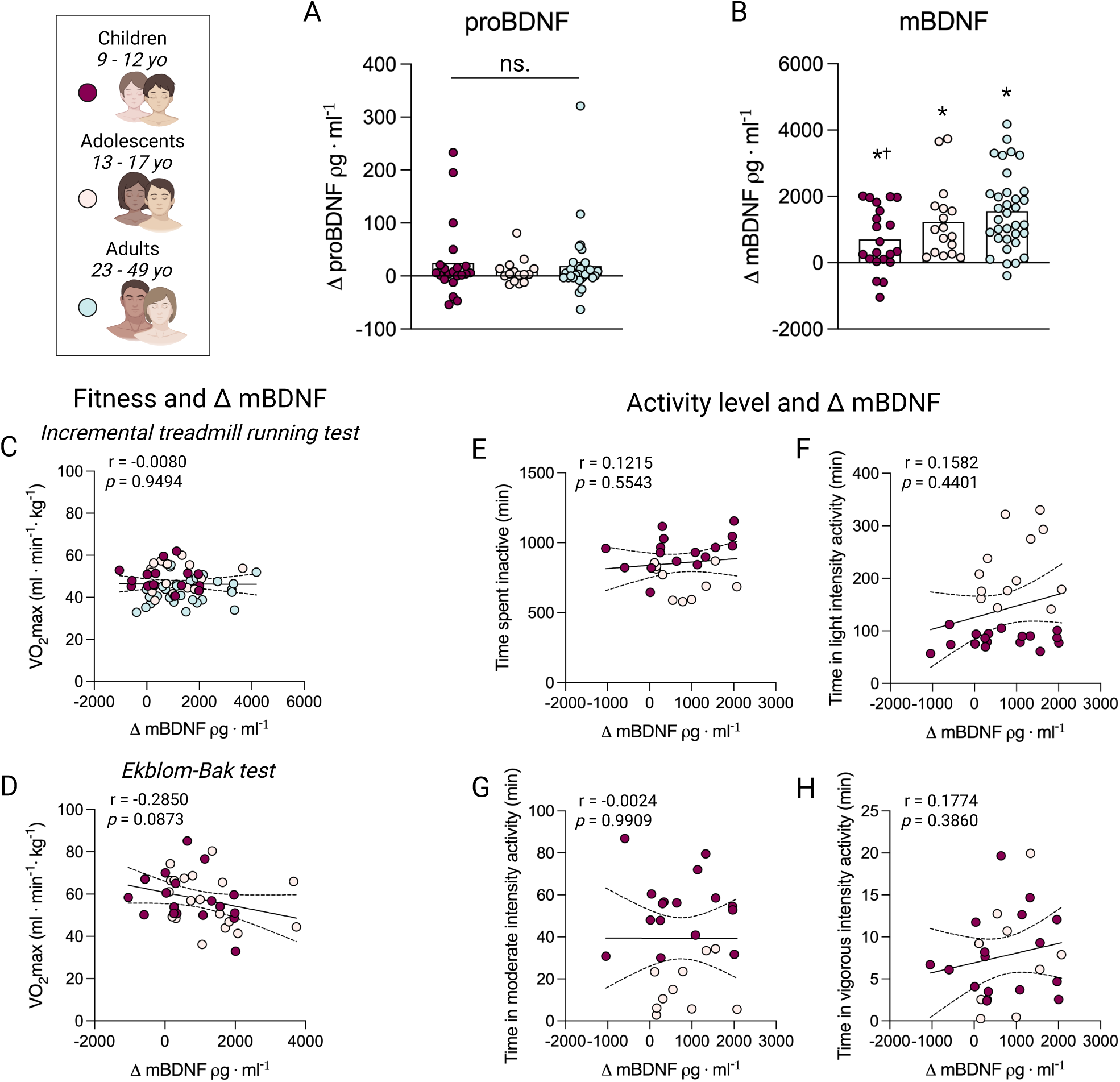
Plasma pro– and mBDNF response to endurance exercise in children, adolescents, and adults. **A-B**: Delta proBDNF levels (A) and, delta mBDNF levels (B) in plasma before and after 45 min running. ns. = not significant. * = p<0.05 vs Pre, † = p<0.05 vs adults. Time effect was analyzed using a two-way ANOVA; Delta values were analyzed with a one-way ANOVA. **C-D**: Correlation of delta plasma mBDNF levels pre and post running with VO_2_max measured during incremental treadmill running (C) and estimated VO_2_max using the Ekblom-Bak test in children and adolescents (D). **E-H**: Correlation of delta plasma mBDNF levels pre and post 45 min running with time spent inactive (E), in light activity (F), in moderate-intensity activity (G), and in vigorous-intensity activity (H) during the day in children and adolescents. C-H was analyzed using Pearson correlation. VO_2_max = Maximal oxygen uptake, Δ = Delta.

Next, we assessed whether the total work performed, or the intensity of the running bout was associated with the acute mBDNF increase in plasma following endurance exercise. Both running distance (Figure 3A), and heart rate intensity (Figure 3B) tended to correlate with the acute mBDNF response to exercise. However, the correlations were weak in both cases (r=0.2127 and r=0.2399, respectively), suggesting a low association. Blood lactate measured at the end of the running bout, as well as the Borg RPE averaged throughout the running bout, showed no association with a change in plasma mBDNF levels (Figures 3C-D).

**Figure 3.**
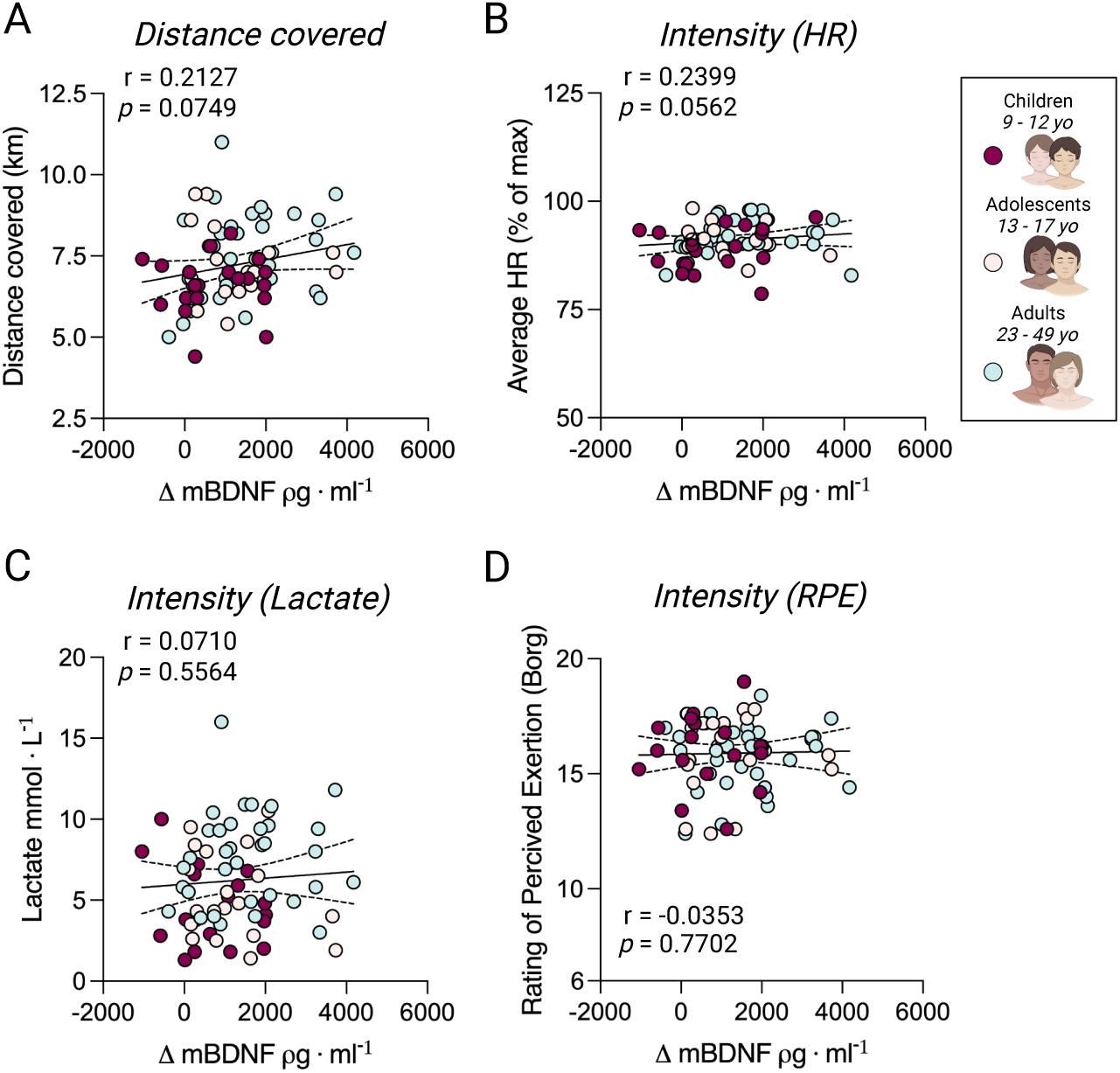
Plasma mBDNF response to endurance exercise and exercise intensity. **A-D**: Correlation of delta mBDNF pre and post 45 min running and running distance in 45 min (A), relative heart rate intensity (B), lactate levels at the end of exercise (C), and average rating of perceived exertion throughout the 45 min run (D). A & C; was analyzed with Pearson correlation, B & D; was analyzed with Spearman correlation. HR = Heart rate, RPE = Rating of Perceived Exertion, Δ = Delta.

Likewise, the same intensity parameters (running distance, heart rate intensity, blood lactate, and RPE) were not associated, or showed a very weak association, with levels of mBDNF in the post-exercise samples (Supplemental Figures 3A-D). Similarly, running distance, heart rate intensity, blood, lactate and RPE were not associated with either delta-(Supplemental Figure 4) or post-exercise proBDNF levels (Supplemental Figure 5).

## Discussion

In this study, we recruited 20 children (9-12 yo), 19 adolescents (13-17 yo), and 39 adults (23-49 yo) to perform a 45-minute endurance exercise session and to donate venous blood samples before and after the exercise. We measured concentrations of pro– and mBDNF in platelet-poor plasma before and after the running bout. The major findings are that: 1) A 45-minute endurance exercise bout elevates plasma mBDNF in youths and adults alike, with adults and the more mature adolescents showing a greater response; 2) intensity during the endurance exercise bout appeared not to play a critical role on the acute mBDNF response; and 3) basal levels of pro– and mBDNF are not associated with age, maturity, or fitness, but rather time spent in vigorous activity.

We show that neither age (Figure 1A-B) nor maturity (Supplemental Figure 1A-F) influences basal plasma levels of pro– and mBDNF in a healthy cohort of participants aged 9-49 years. This is in contrast to previous findings describing that pubertal males had significantly lower plasma BDNF levels, as compared to prepubertal boys and girls^39^. Although our data indicate numerically lower plasma mBDNF levels in adolescents (13-17 yo) compared to children (9-12 yo; Figure 1B), this was not mediated by lower values in the adolescent (data not shown) as suggested by previous observations^39^. Our data on mBDNF levels and maturity (Tanner stages & circulating sex hormones; Supplemental figure 2) also support an opposite association, where more mature children seem to have slightly elevated basal plasma mBDNF levels. We also suspect that the numerically lower mBDNF values observed in adolescents compared to children in our data may be related to daily physical activity (Figure 1E-H). Specifically, the adolescents, as compared to the children, spent more time doing light-intensity activity at the expense of moderate-intensity activity, which may be more influential over basal plasma mBDNF levels, although the relationship was very weak (Figure 1F-G).

Our data reveal a significant positive correlation between time spent in vigorous physical activity and basal plasma mBDNF levels (r=0.5794, p=0.0015; Figure 1H) in children and adolescents. In contrast, no significant associations were observed with time spent inactive or in light-or moderate-intensity activity (Figure 1E-G). The exclusive correlation with high-intensity activity aligns with the notion that circulating BDNF levels may be modulated by exercise intensity, as suggested in previous studies^18^. However, the literature on this topic remains inconsistent, with some studies reporting a negative association between physical activity and serum BDNF levels^40^, while others, using plasma mBDNF, report either no association^24^ or a positive relationship^23^. Interestingly, while time spent in vigorous-intensity activity was a predictor of VO_2_max in children and adolescents (Figure 1L), there was no correlation between fitness levels and basal plasma mBDNF (Figure 1C-D) or proBDNF (data not shown). These findings collectively suggest that basal plasma mBDNF levels may be more reflective of the cumulative effects of repeated exercise-induced “mBDNF release pulses” rather than an individual’s overall physical fitness.

To the best of our knowledge, this study is the first to examine the plasma pro– and mBDNF response to acute exercise in both children and adolescents. Previous studies have primarily focused on basal serum mBDNF levels in response to long-term exercise programs^41–43^, leaving a gap in our understanding of the acute exercise-induced changes in circulating BDNF in younger populations. Our findings demonstrate that plasma mBDNF levels increased across all groups—children, adolescents, and adults—following a 45-minute running bout. Notably, children exhibited a significantly lower mBDNF response compared to adults (Figure 2B), while the response in adolescents was intermediate, with a numerical increase greater than that of children but less than that of adults. This gradient effect, where the youngest participants showed the smallest increase and adults showed the largest, suggests an age-related difference in the acute mBDNF response to endurance exercise.

Further supporting an age or maturity effect, children in more advanced Tanner stages (4 and 5) exhibited a significantly greater mBDNF response compared to those in Tanner stages 2 and 3 (p<0.05; Supplemental Figure 2). This suggests that biological maturity plays a role in modulating the BDNF response to exercise. Additionally, we observed a near-significant association between estradiol levels and the change in mBDNF in boys (r=0.391, p=0.088), while no such association was detected in girls (r=0.264, p=0.307). The absence of a correlation in girls may be attributed to the monthly hormonal fluctuations that were not accounted for in some of the female post-pubertal participants. Interestingly, testosterone levels showed a borderline significant association with mBDNF changes in both boys (r=0.397, p=0.083) and girls (r=0.357, p=0.159). However, given the strong correlation between sex hormones (data not shown), it is possible that the observed relationship with testosterone is secondary to the influence of estradiol.

Contrary to our observations regarding basal plasma mBDNF levels, the acute mBDNF response to exercise did not correlate with physical activity levels (Figure 2E-H).

Additionally, we found that underlying fitness levels did not significantly influence the exercise-induced increase in plasma mBDNF (Figure 2C-D). This lack of association persisted even when the analysis was restricted to adults (data not shown), which contrasts with previous reports suggesting that higher fitness levels^20^ or prior exercise training^19^ may enhance the acute mBDNF response to endurance exercise.

We did not observe an increase in plasma proBDNF immediately following the endurance exercise session (Figure 2A), which contrasts with our previous findings where resistance exercise led to a modest rise in proBDNF levels^5^. This was unexpected, given that proBDNF, unlike mBDNF, is predominantly associated with slow-twitch type I muscle fibers^5^. Since endurance exercise relies more heavily on type I fiber activity, a similar or even greater release of proBDNF into circulation following the endurance session could be expected. One possible explanation for the absence of a detectable increase in proBDNF is that the running may have triggered the release of proBDNF earlier in the exercise session. As our blood samples were collected only at the conclusion of the 45-minute session, it is possible that proBDNF had already been released and cleaved into mBDNF by this time, reducing its detectable levels in the plasma^4^. Additionally, the self-paced nature of the running protocol could have contributed to variable proBDNF release among participants. Some participants began the session at a high intensity, while others adopted a more cautious approach, as reflected in their heart rate data. This variability may have resulted in a more varied release of proBDNF in our current cohort, in contrast to the more uniform resistance exercise protocol used in our previous study^5^. Finally, as mBDNF release seems to be limited to vigorous-intensity activity (Figure 1E-H), it may be that proBDNF release from skeletal muscle also requires an unknown intensity threshold or an additional mechanical loading component to stimulate its release, which would only be attainable from resistance exercise.

Our finding that exercise intensity did not correlate with the proBDNF and mBDNF response to the exercise session was unexpected (Figure 3 & Supplemental Figure 3), particularly given previous studies suggesting that mBDNF release into circulation is intensity-dependent^18^. Although both total running distance (Figure 3A) and heart rate intensity (Figure 3B) approached significance, with p-values of 0.0749 and 0.0562, respectively, the correlations were weak, with R-values close to 0.2. This lack of an association between changes in mBDNF and exercise intensity might be explained by the study design, in which participants were instructed to run at a pace that would allow them to maximize distance within the 45-minute session. It is also possible that, given the high exercise intensity (91 ± 4% of maximum heart rate; Figure 3B), all participants were already exercising at a level sufficient to trigger maximal mBDNF release. However, the suggested notion that mBDNF release may be intensity-dependent is primarily based on measurements in serum^18^, where mBDNF concentrations can be up to 100-fold higher than in platelet-poor plasma^6,44^. Thus, it may be that the observed association between exercise intensity and mBDNF release is influenced by the exercise-mediated release of platelets, which has also been reported to be intensity-dependent^45^.

In our previous work, we observed an increase in plasma mBDNF levels in response to lactate infusion during resistance exercise^5^, suggesting that further increases in exercise intensity, and consequently blood lactate levels, could potentially enhance mBDNF release. However, in the present study, we found no significant correlation between post-exercise blood lactate levels and either the change in mBDNF levels or the absolute mBDNF levels after the 45-minute run (Figure 3B & Supplemental Figure 3B). Given the elevated blood lactate (6.3 ± 3.0 mmol × L⁻¹) and mBDNF levels (2216 ± 1400 pg × ml⁻¹) observed here, it is possible that the effect of lactate on mBDNF release had already reached saturation at this high exercise intensity. This would align with findings from murine models, where higher lactate injections (180 mg × kg⁻¹) did not further increase mBDNF production beyond the levels observed with smaller doses (117 mg × kg⁻¹), indicating a possible plateau effect^46^.

Further, it is possible that a lactate-stimulated increase in BDNF production within the hippocampus^46^ and a potential subsequent release into the circulation may be small and thus hard to distinguish in a self-paced exercise session such as the one used in the current intervention.

This study demonstrates that plasma mBDNF levels increase following endurance exercise across children, adolescents, and adults, with a more pronounced response observed in adults and more mature adolescents. Basal mBDNF levels were associated with time spent in vigorous physical activity, while the acute exercise response was not significantly influenced by exercise intensity or baseline fitness levels. These findings highlight the importance of biological maturity in modulating the neurotrophic response to exercise and contribute to our understanding of how BDNF responds to physical activity across childhood, adolescence, and adulthood.

## Acknowledgments

The authors want to thank nurses Emelie Flodin and Björn ‘Bloody Bear’ Lanzky Otto for excellent assistance with blood sampling throughout the study. The authors would also like to thank the Swedish Sports Confederation’s development center, Bosön, Stockholm, Sweden, for kindly providing facilities for indoor running. Moreover, the authors would like to thank the Department of Biobank and Study Support at Karolinska University Hospital for their contribution, including professional service and support. Figures were created with Biorender and GraphPad.

## Author Contributions

Writing – Original draft: SE, JS

Writing – Reviewing and editing: All authors

Conceptualization: SE, LC, MM, FvW

Data collection: SE, JS, LC, WH, AvF, SR, JN, FvW

Data analysis: SE, JS, LC, WH, MR, MM

Data interpretation: SE, JS, MM, FvW

Data visualization: SE

Funding acquisition: SE, JN, MM, FvW

## Consent Statement

All participants were informed orally and in writing as to the risks and benefits of enrollment, and informed consents were obtained from all participants, as well as from the caregivers of children under the age of 15.

---

**Supplemental Figure 1.**
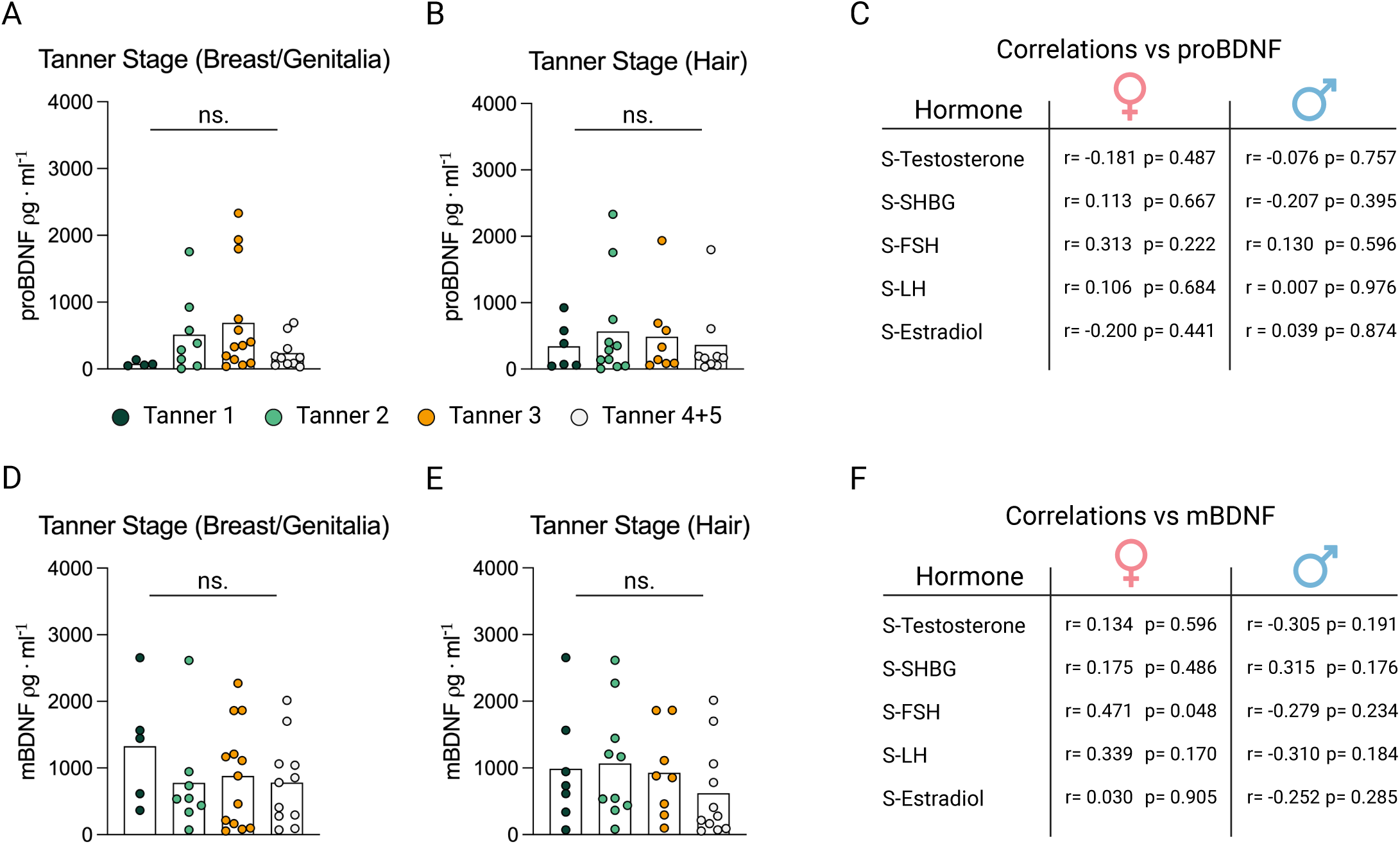
Maturity in children and adolescents and basal plasma pro– and mBDNF levels. **A-B**: Tanner stage assessment and plasma proBDNF levels at rest in children and adolescents. **C**: Correlations of plasma proBDNF levels at rest and sex hormone concentrations in blood. **D-E**: Tanner stage assessment and plasma mBDNF levels at rest in children and adolescents. **F**: Correlations of plasma mBDNF levels at rest and sex hormone concentrations in blood. A, B, D & E were analyzed with a one-way ANOVA, ns = not significant. C & F were analyzed with Pearson correlation. SHGB = Sex hormone binding globulin, FSH = Follicle stimulating hormone, LH = Luteal hormone.

**Supplemental Figure 2.**
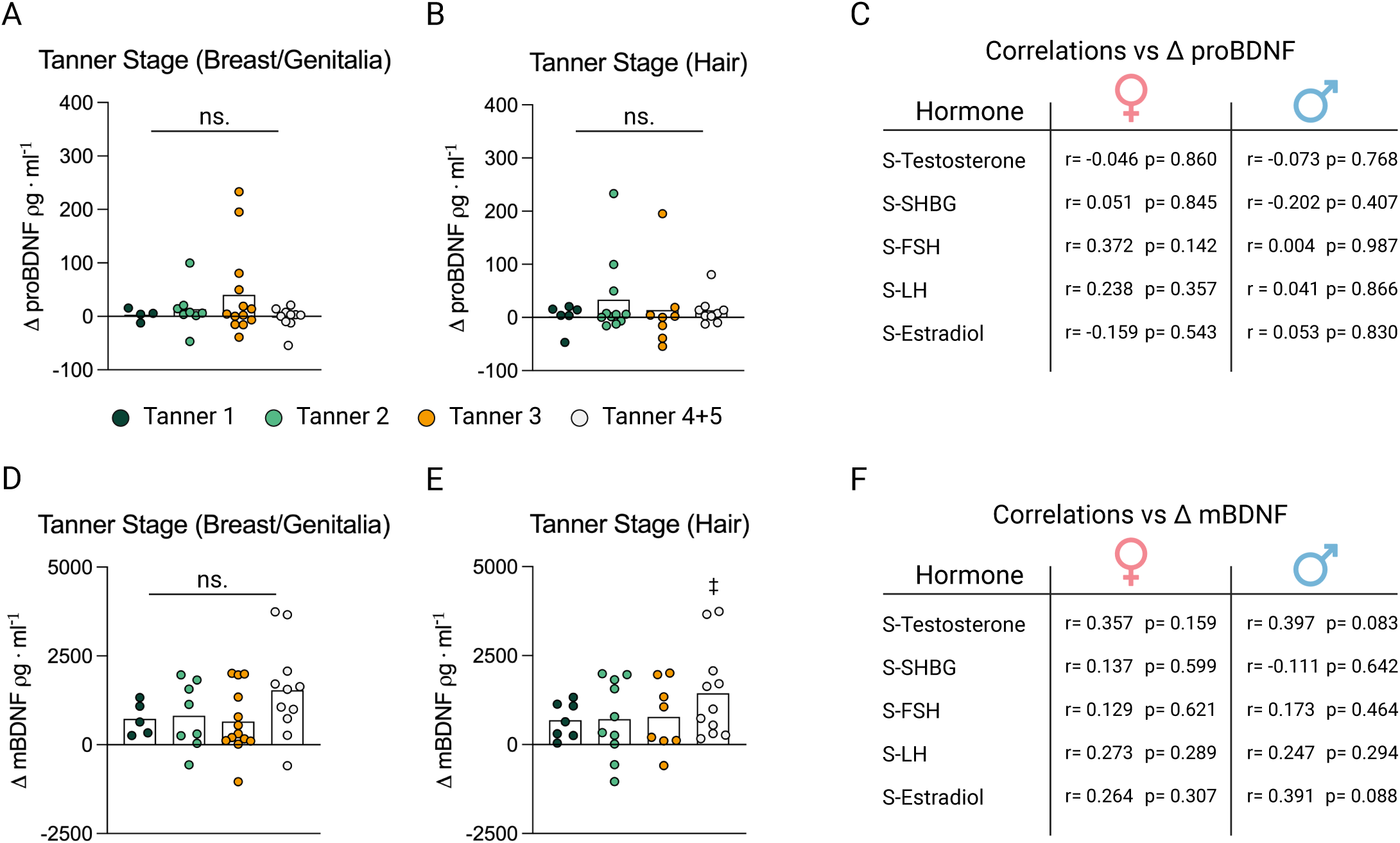
Maturity in children and adolescents and plasma pro– and mBDNF response to endurance exercise. **A-B**: Tanner stage assessment and delta plasma proBDNF levels pre and post a 45 min run in children and adolescents. **C**: Correlations of delta plasma proBDNF levels pre and post a 45 min run and sex hormone concentrations in blood. **D-E**: Tanner stage assessment and delta plasma mBDNF levels pre and post a 45 min run in children and adolescents. **F**: Correlations delta plasma mBDNF levels pre and post a 45 min run and sex hormone concentrations in blood. A, B, D & E were analyzed with a one-way ANOVA, ns = not significant, ‡ = p< 0.05 vs Tanner 2&3. C & F were analyzed with Pearson correlation. SHGB = Sex hormone binding globulin, FSH = Follicle stimulating hormone, LH = Luteal hormone, Δ = Delta.

**Supplemental Figure 3.**
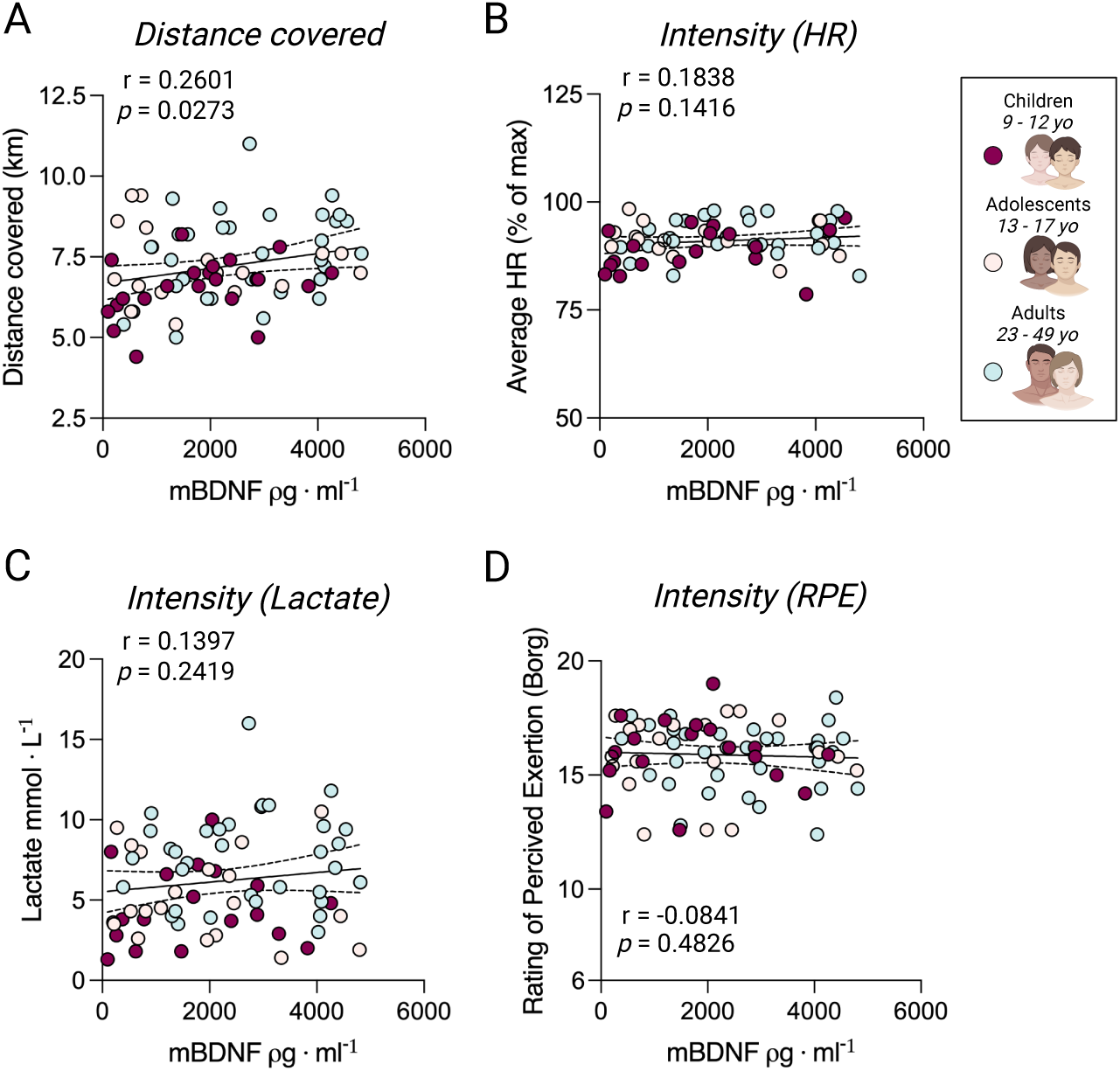
Plasma mBDNF post endurance exercise and exercise intensity. **A-D**: Correlation of mBDNF post 45 min running with running distance in 45 min (A), relative heart rate intensity (B), lactate levels at the end of exercise (C), and average rating of perceived exertion throughout the 45 min run (D). A & C; A & C; was analyzed with Pearson correlation, B & D; was analyzed with Spearman correlation. HR = Heart rate, RPE = Rating of Perceived Exertion, Δ = Delta.

**Supplemental Figure 4.**
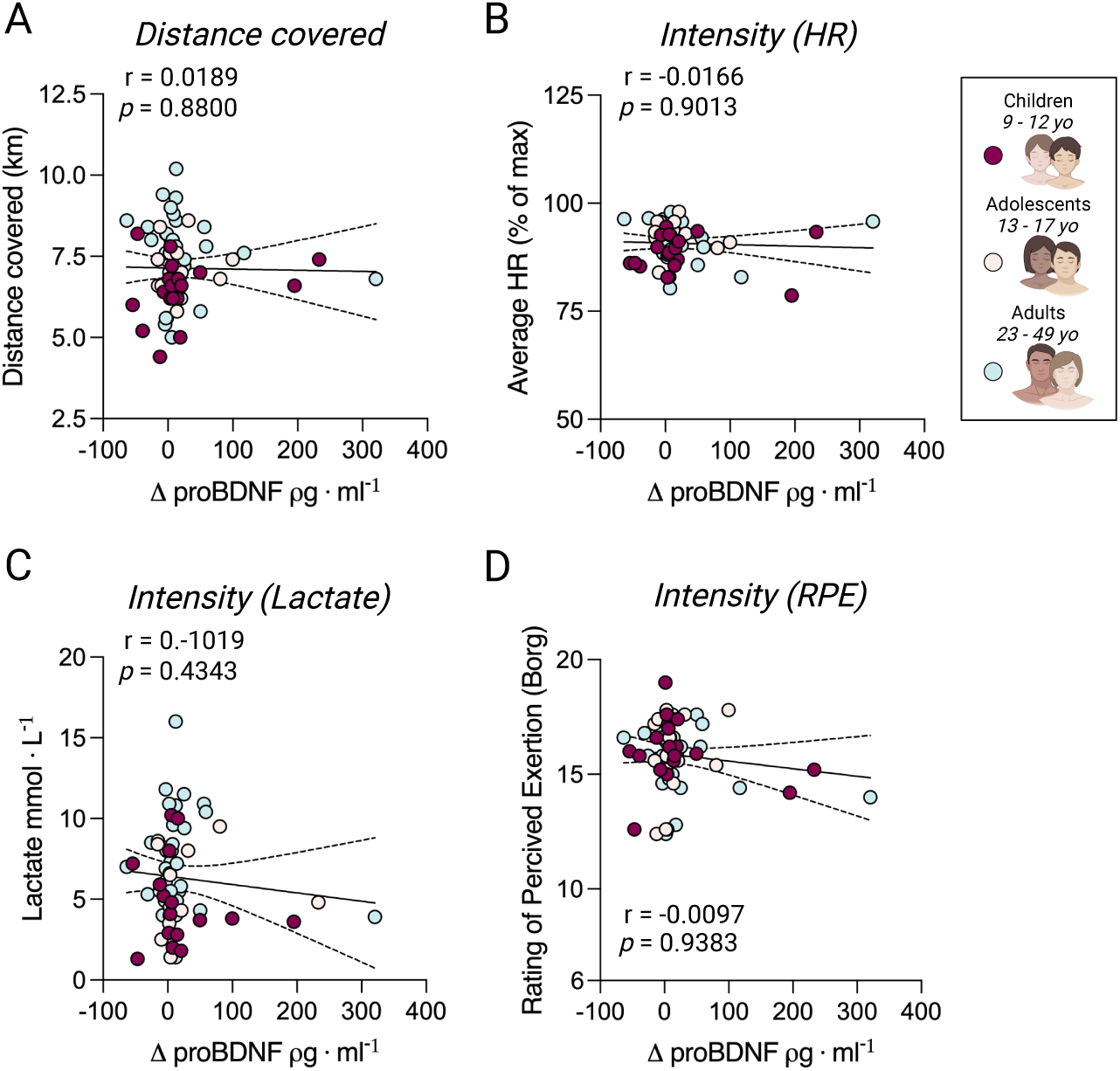
Plasma proBDNF response to endurance exercise and exercise intensity. **A-D**: Correlation of delta proBDNF pre and post 45 min running and running distance in 45 min (A), relative heart rate intensity (B), lactate levels at the end of exercise (C), and average rating of perceived exertion throughout the 45 min run (D). A & C; was analyzed with Pearson correlation, B & D; was analyzed with Spearman correlation. HR = Heart rate, RPE = Rating of Perceived Exertion, Δ = Delta.

**Supplemental Figure 5.**
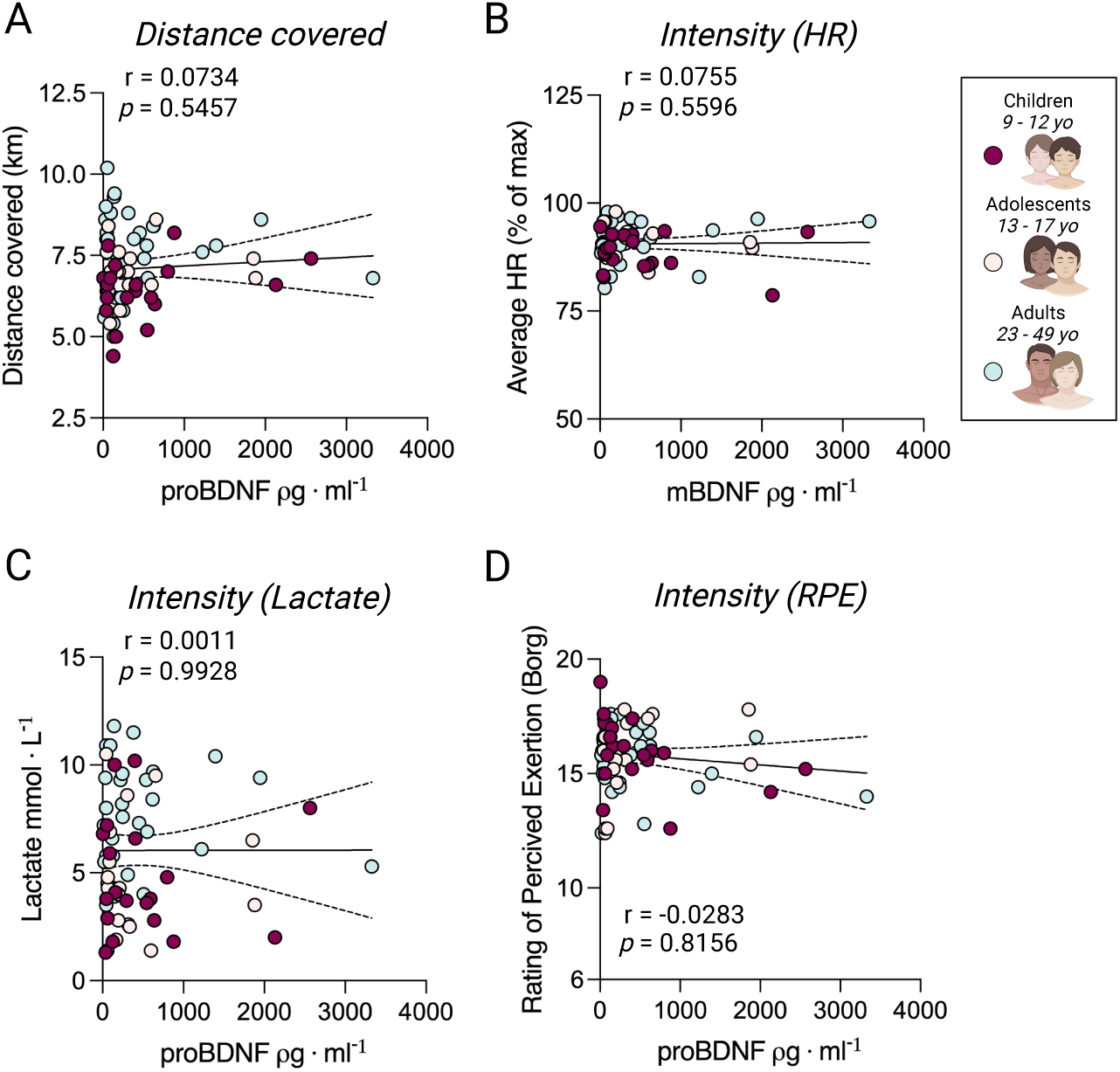
Plasma proBDNF post endurance exercise and exercise intensity. **A-D**: Correlation of proBDNF post 45 min running with running distance in 45 min (A), relative heart rate intensity (B), lactate levels at the end of exercise (C), and average rating of perceived exertion throughout the 45 min run (D). A & C; A & C; was analyzed with Pearson correlation, B & D; was analyzed with Spearman correlation. HR = Heart rate, RPE = Rating of Perceived Exertion, Δ = Delta.

